# The TRIP6/LATS1 complex constitutes the tension sensor of α-catenin/vinculin at both bicellular and tricellular junctions

**DOI:** 10.1101/2023.06.05.543720

**Authors:** Lin Xie, Gangyun Wu, Xiayu Liu, Xiufen Duan, Kaiyao Zhou, Hua Li, Wenxiu Ning

**Author notes:** Corresponding authors: Wenxiu Ning; Hua Li.

## Abstract

Cell-cell mechanotransduction regulates tissue development and homeostasis. α-catenin, the core component of adherens junctions, functions as a tension sensor and transducer by recruiting vinculin and transducing the signals to influence cell behaviors. However, little is known about the components, distributions, and dynamics of the α-catenin based tension sensors at the cell junctions. Here, we uncovered the TRIP6/LATS1 complex locates at the tension sites where α-catenin/vinculin is at both the bicellular junctions (BCJs) and tricellular junctions (TCJs). Vinculin/TRIP6/LATS1 are prone to form as puncta in the cytoplasm without α-catenin participation. Furthermore, the tension sensing complex distributed stronger at TCJs and exhibited a discontinuously button-like pattern on BCJs. The α-catenin/vinculin BiFC-based mechanosensor further proved the discontinuous distribution of the tension at BCJs, and was more motile than the TCJs. In summary, our study revealed that TRIP6 and LATS1 are novel compositions of the tension sensor, together with the core complex of α-catenin/vinculin, at both the BCJs and TCJs. This work gives insights and improvements in exploring the molecular mechanism that mediates cell-cell mechanotransduction at cell junctions.

## Introduction

Cells in tissues perceive and integrate the mechanical signals, which include tension, compression and shear stress. These signals are essential for maintaining tissue structure, function, and homeostasis (Hayward et al., 2021). Cells sense the forces and transduce the mechanical cues into biochemical signals to stimulate a process called mechanotransduction (Broussard et al., 2020; Saraswathibhatla et al., 2023). The mechanical cues are sensed by several stucutres including mechanosensitive ion channels (Brohawn et al., 2014; Murthy et al., 2018; Pan et al., 2018; Saotome et al., 2018), integrins, and the adherens junctions (AJs) mediated by cadherins (Gauthier and Roca-Cusachs, 2018; Horne-Badovinac, 2014), that transduce the signals to remodel the cytoskeleton and modulates gene expression. Mechanotransduction ultimately influences cellular behaviors such as cell adhesion, migration, proliferation, differentiation, and survival (Dogterom and Koenderink, 2019; Mui et al., 2016). Abnormal stress of cell mechanotransduction is associated with developmental defects, inflammatory diseases, and tumor growth and metastasis (Gomez-Gonzalez et al., 2020; Sugimura et al., 2016).

AJs are essential structures that mediate cell-cell mechanotransduction (Angulo-Urarte et al., 2020). The cadherin–catenin complex, consisting of E-cadherin, α-catenin and β-catenin, plays a critical role in mediating epithelial cell–cell interactions. α-catenin acts as the tension sensor and transducer at AJs, which is closely related to its force-sensitive conformation and interactions with vinculin. In response to high tension, α-catenin opens its VH2 domain for vinculin binding. The bond vinculin turns into an open conformation, and recruit Mena/VASP, ultimately assembling F-actin in a force-dependent manner (Drees et al., 2005; Mege and Ishiyama, 2017). Therefore, α-catenin/vinculin is considered as the core complex of AJs that senses and responds to mechanical forces. The α-catenin/vinculin complex recruits TRIP6 to the tension site, which further inhibits LATS1 and inhibits its activity while activating YAP (Adhikari et al., 2018; Dutta et al., 2018; Thomas et al., 2013). Tricellular junctions (TCJs) are areas where mechanical force is applied to cell adhesion molecules and structures, and experience high tension due to the tensile forces along three BCJs that are applied around cell vertices (Bosveld et al., 2018; Higashi and Miller, 2017; Trichas et al., 2012). It was recently reported that α-catenin acts as a new binding partner of tricellulin at TCJs, and bridging tricellulin attachment to the bicellular actin cables to support the epithelial barrier at cell vertices (Cho et al., 2022). Vinculin is also recruited to TCJs similar to BCJs in response to tension which is dependent on α-catenin as well (Cho et al., 2022). Although integrin-mediated cell-matrix derived mechanotransduction has been well-illustrated, studies on cell-cell mechanotransduction mediated by the α-catenin/vinculin tension sensor at bicellular junctions (BCJs) as well as TCJs are still limited.

In this study, by using split-TurboID, mutations of α-catenin, as well as BiFC-based mechanosensor, we proved that TRIP6/LATS1 complex is located not only at BCJs but also at TCJs where the α-catenin/vinculin interaction is. The TRIP6/LATS1 complex forms as condensates with vinculin but not α-catenin in the cytoplasm. Moreover, we found that the TRIP6/LATS1/α-catenin/vinculin cassette was found to have a discontinuously button-like distribution at BCJs and higher intensity at TCJs. Finally, the BiFC mechanosensor based on the interactions between α-catenin and vinculin further supported this observation and also demonstrated the dynamics of the tension sensor at BCJs and TCJs.

## Results and Discussion

### Identification of LATS1 as a component of the tension sensor using split-TurboID between α-catenin and vinculin

To explore possible factors that might involve in AJs-mediated cell-cell mechanotransduction, we took advantage of the force-responsive interactions between α-catenin and vinculin at AJs. Split-TurboID which could label proximal proteins of two interactors was applied to detect factors that are proximal to α-catenin/vinculin complex (Cho et al., 2020) (Figure 1A). We employed an IRES sequence dependent second gene expression system to co-express Flag-α-catenin-spTurboN and spTurboC-vinculin-HA from a single promoter. The constructed all-in-one plasmid was called Flag-α-catenin-spTurboN-IRES-spTurboC-vinculin-HA, which was designed to detect the interactome of the tension sensing complex of α-catenin/vinculin (Figure 1B), hereafter named mechano-spTurboID. Both the expression and protein sizes of Flag-α-catenin-spTurboN and spTurboC-vinculin-HA were confirmed right by the immunofluorescence staining and western blotting (Figures 1C-D).

**Figure 1.**
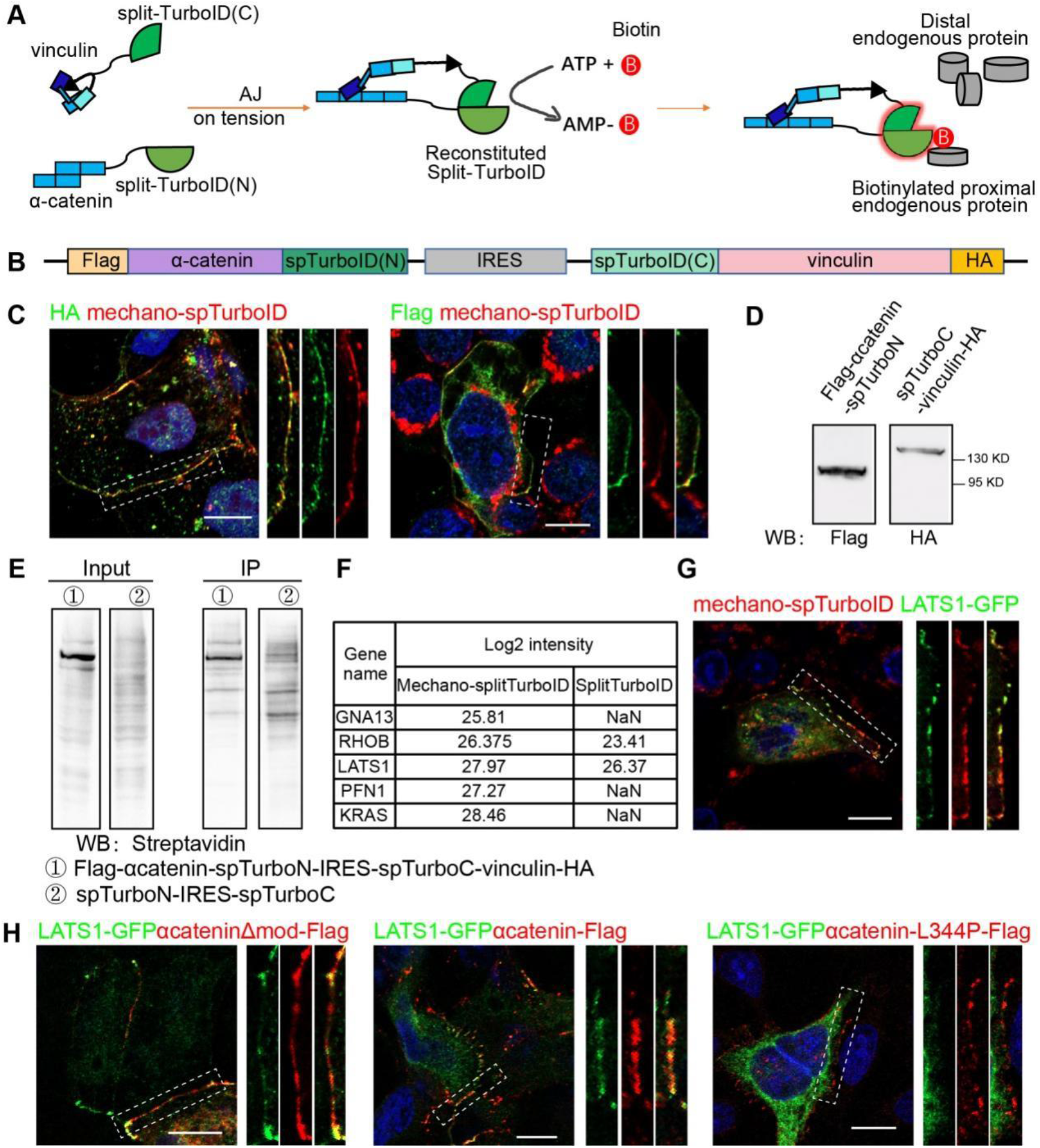
Split-TurboID-based proteomics identified LATS1 co-localized with the tension sensor at cell junctions. **A**. Mechano-splitTurboID model based on the tension dependent interaction between α-catenin and vinculin. **B**. All-in-one plasmid design of mechano-spTurboID based on α-catenin and vinculin interaction. **C**. Immunofluorescence staining of streptavidin-labeled biotinylated proteins (red) and spTurboC-vinculin-HA (left), Flag-α-catenin-spTurboN (right) in mechano-spTurboID transfected DLD1 cells, respectively. **D**. Western blotting confirmation of Flag-α-catenin-spTurboN and spTurboC-vinculin-HA expression in transfected HEK293T cells. **E**. Western blotting of the streptavidin-labeled biotinylated proteins in mechano-spTurboID and spTurboID control transfected cells. **F**. List of selected candidate genes enriched in the mechano-spTurboID mass spectrometry dataset. **G**. Immunofluorescence staining of LATS1-GFP (green) and streptavidin-labeled biotinylated proteins (red) in the mechano-spTurboID transfected DLD1 cells. **H**. Immunofluorescence staining of LATS1-GFP (green) with α-catenin-Δmod-Flag (left), α-catenin-Flag (middle), and α-catenin-L344P-Flag (right) in DLD1 cells. All dashed box regions were enlarged on the right. Scale bars are all 10 μm.

We further purified the biotinylated proteins in mechano-spTurboID transfected 293T cells using streptavidin conjugated beads. The purified biotinylated proteins in mechano-spTurboID showed weak but different bands compared with the split-TurboID control confirmed by western blotting (Figure 1E). The purified biotinylated proteins were further analyzed using label-free quantitative mass spectrometry. LATS1, a YAP1 upstream regulator (Totaro et al., 2018), was enriched in the mechano-spTurboID interactome compared with the control (Figure 1F). LATS1 has been reported to localize at AJs in response to tension dependent on TRIP6 (Dutta et al., 2018). To confirm LATS1 junctional localization during cell-cell mechanotransduction, we constructed LATS1-GFP plasmid and co-transfected it with mechano-spTurboID in DLD1 epithelial cells. It showed LATS1 had a strong colocalization with the mechano-spTurboID with discontinuous localization signals at the BCJs (Figure 1G).

Depletion of the Mod domain abolished the self-inhibition of α-catenin, which could expose vinculin binding sites and constitutively recruit vinculin, while mutation of L344P could abolish its interaction with vinculin (Figures S1A) (Le et al., 2021). Co-staining of full-length α-catenin, α-catenin mutants of Δmod-Flag and L344P-Flag confirmed the enhanced interaction of vinculin and α-catenin in Δmod-Flag but not L344P-Flag, compared with α-catenin-Flag (Figures S1B-D). Notably, vinculin-HA colocalization with Δmod-Flag also showed a discontinuous distribution at AJs. Co-staining of full-length α-catenin, Δmod-Flag, and L344P-Flag with LATS1 further confirmed the colocalization of LATS1 with α-catenin/vinculin complex at BCJs in a discontinuous distributed pattern (Figure 1H).

LATS1 is a core kinase in the Hippo pathway that inhibits YAP through phosphorylation (Gan et al., 2020; Strano and Blandino, 2007). Phosphorylated YAP further interacts with 14-3-3 and thus gets detained at the cytoplasm (Meng et al., 2016). Researches showed that α-catenin can interact with YAP and also 14-3-3 (Schlegelmilch et al., 2011). Therefore, we further examined the possible relationship of YAP (Figures S2A-C) and 14-3-3 (Figures S2D-F) with the complex of α-catenin and vinculin at BCJs. Staining of YAP1 and 14-3-3 with full-length α-catenin, Δmod-Flag, and L344P-Flag showed no obvious colocalization. Thus, LATS1 is the main component in the Hippo pathway that responds to the AJs based mechanotransduction.

### The TRIP6/LATS1 complex colocalizes with the tension sensor of α-catenin/vinculin at BCJs and is distributed discontinuously

Notably, TRIP6 is reported to interact with LATS1/2 kinases by competing with MOB1 and recruits LATS1/2 to junctions under tension (Dutta et al., 2018; Venkatramanan et al., 2021). We then tested the relationship of TRIP6 with the α-catenin/vinculin/LATS1 complex at the tension sites. Co-staining of TRIP6-GFP with full-length α-catenin, Δmod-Flag, and L344P-Flag confirmed TRIP6 colocalized with α-catenin/vinculin complex at BCJs (Figures 2A-C). Colocalization of the TRIP6 with α-catenin/vinculin/LATS1 complex at BCJs were further confirmed (Figures 2D-F). Thus, the results confirmed that TRIP6 and LATS1 complex appears at AJs where α-catenin and vinculin interacts. Interestingly, this α-catenin/vinculin/TRIP6/Lats1 cassette, Δmod-Flag, as well as the initial mechano-TurboID all showed discontinuous distribution at AJs. As the interaction of α-catenin and vinculin occurs in response to the high tension, the discontinuous distribution of this cassette implicates the unevenly distributed forces exerting on BCJs.

**Figure 2.**
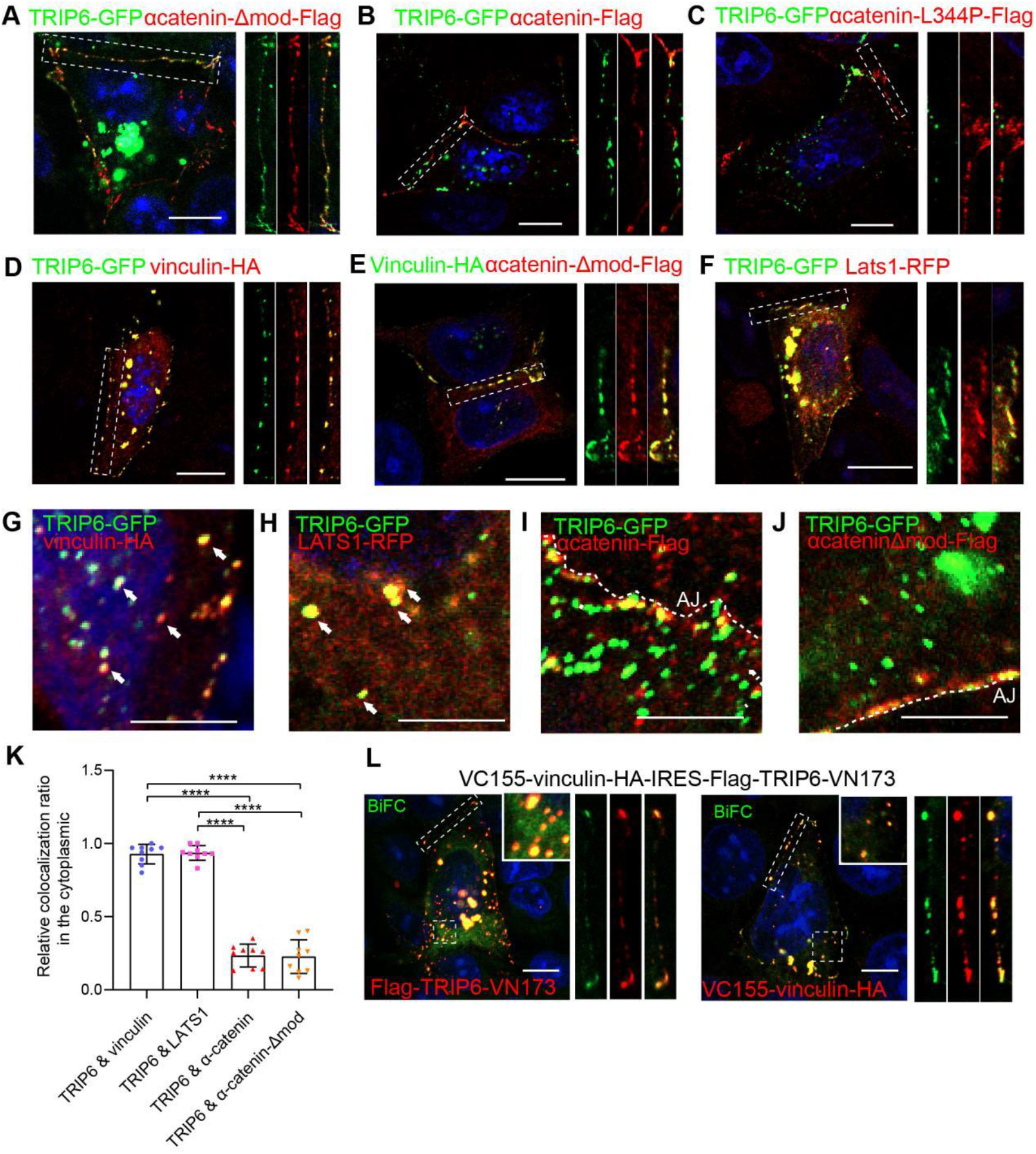
TRIP6/LATS1/Vinculin are prone to form puncta in the cytoplasm and co-localize with junctional α-catenin. **A-D**. Immunofluorescence staining of TRIP6-GFP (green) with α-cateninΔmod-Flag (A), α-catenin-Flag (B), α-catenin-L344P-Flag (C), and vinculin-HA (D) in DLD1 cells, respectively. **E**. Immunofluorescence staining of vinculin-HA (green) and α-catenin-Δmod-Flag (red) in DLD1 cells. **F**. Immunofluorescence staining of TRIP6-GFP (green) and LATS1-RFP (red) in DLD1 cells. **G-J**. Immunofluorescence staining of TRIP6-GFP (green) with vinculin-HA (G), LATS1-RFP (H), α-catenin-Flag (I), and α-catenin-Δmod-Flag (J) in DLD1 cells, respectively. Arrows indicate colocalized puncta in the cytoplasm. Dashed lines indicate the AJs. **K**. Colocalization ratio of the cytoplasmic puncta in different conditions. N=9 cells from each group. Data were analyzed by one-way ANOVA using Tukey’s multiple comparisons test. **** means p value <0.0001. **L**. The BiFC signal (green) in VC155-vinculin-HA-IRES-Flag-TRIP6-VN173 transfected DLD1 cells, and VC155-vinculin-HA (left) or Flag-TRIP6-VN173 (right) was shown in red. All dashed box regions were enlarged on the right. Scale bars in G-J are 5 μm, the rest ones are 10 μm.

The TRIP6-GFP also exhibited as puncta distribution in the cytoplasm and these puncta appeared unrelated with α-catenin or Δmod-Flag, while showing strong colocalization with vinculin-HA and LATS1-RFP (Figures 2G-K). Interestingly, vinculin and LATS1 were prone to form the cytoplasmic puncta only when they were co-transfected with TRIP6-GFP but not with α-catenin or Δmod-Flag (Figures 2D-F, 1H). BiFC, a bimolecular fluorescence complementation assay, commonly used to directly visualize protein-protein interaction (Hu et al., 2002). BiFC assay of TRIP6 and vinculin further confirmed their interaction both in the cytoplasm as puncta and also at the BCJs (Figures 2L).

### The TRIP6/LATS1 complex colocalizes with the tension sensor of α-catenin/vinculin at TCJs

Interestingly, we also observed the strong signal of α-catenin/vinculin/TRIP6/LATS1 cassette, Δmod-Flag, as well as the mechano-TurboID at TCJs (Figures 3A-D). TCJs are specialized cell junctions and formed at sites where three cells contact. We therefore cloned tricellulin, a tight junction protein that is localized in TCJs (Cho et al., 2022). Immunostaining data showed vinculin-HA, Δmod-Flag, and TRIP6-GFP colocalized with tricellulin-RFP (Figures 3E-G). Staining of endogenous LATS1, α-catenin, and TRIP6 with tricellulin-RFP further confirmed the colocalization of α-catenin/vinculin/TRIP6/LATS1 cassette with tricellulin at TCJs (Figures 3H-J). The time-lapse assay also proved the stable localization of TRIP6/LATS1 at TCJs (Movies S1 and S2).

**Figure 3.**
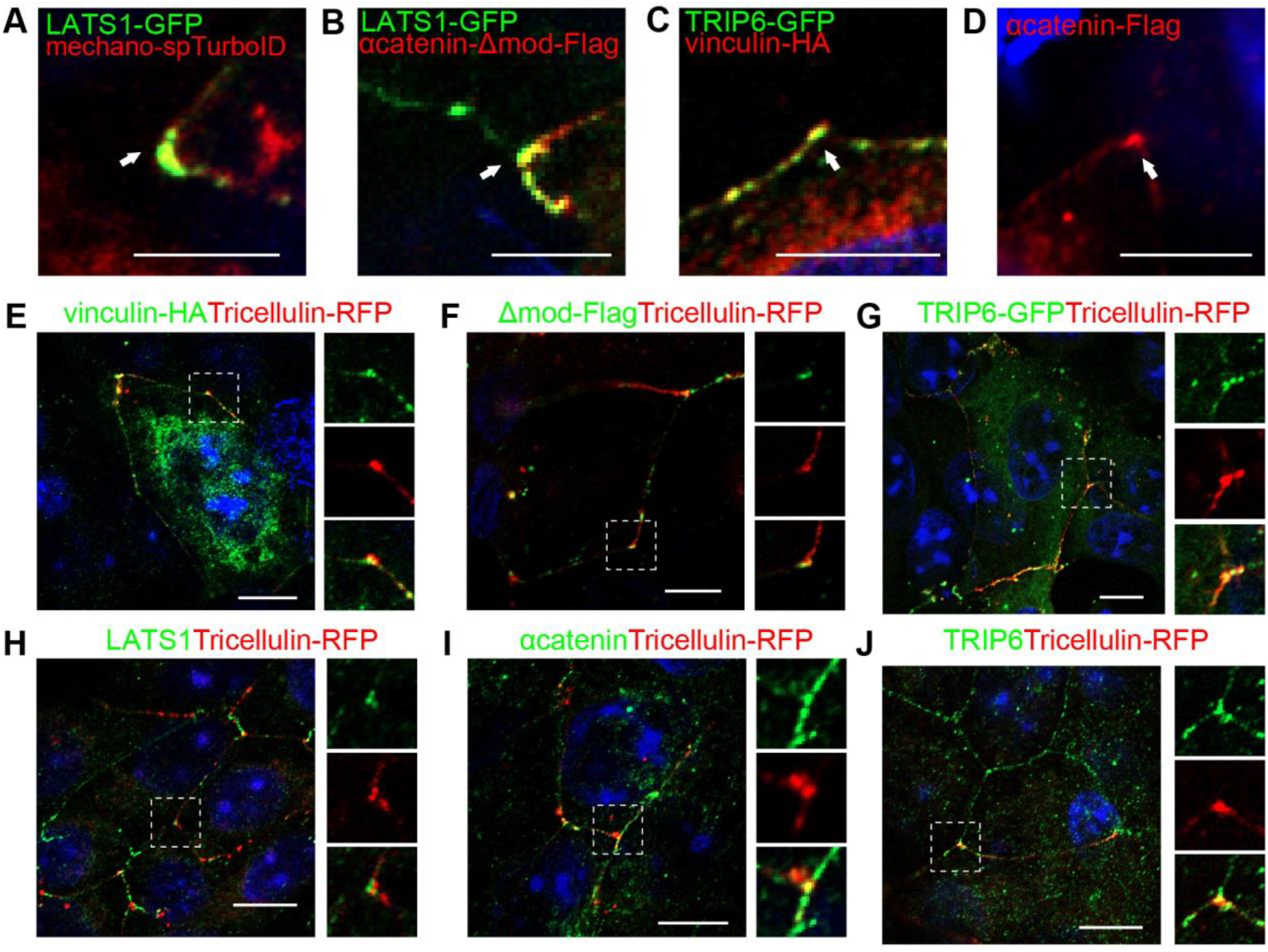
The α-catenin/vinculin/TRIP6/LATS1 complex colocalizes with tricellulin at TCJs. **A-B**. Immunofluorescence staining of LAT1-GFP and the mechano-spTurboID (A) andα-catenin-Δmod-Flag (B) at TCJs. **C**. Immunofluorescence staining of TRIP6-GFP and vinculin-HA at TCJs. **D**. Immunofluorescence staining of α-catenin-Flag at the TCJs. **E**-**J**. Immunofluorescence staining of vinculin-HA (E), α-catenin-Δmod-Flag (F), TRIP6-GFP (G), endogenous LATS1 (H), αcatenin (I), TRIP6 (J) with tricellulin-RFP at the TCJs respectively. All dashed box regions were enlarged on the right. Scale bars in A-D are 5 μm, in E-G are 10 μm.

### A BiFC-based mechanosensor to monitor the dynamics of the tension

An obstacle to investigate cell-cell mechanotransduction is lacking visible sensors to detect and display the spatial and temporal dynamics of the forces. Thus, we designed a mechanosensor based on the interaction of α-catenin and vinculin at the tension sites using the BiFC method (Figure 4A). α-catenin was linked to VN173 plasmid and vinculin was linked to VC155 plasmid, then these two parts were cloned into an all-in-one plasmid by an IRES sequence (Figure 4B). Immunostaining results showed the BiFC signals of this mechanosensor colocalized both with α-catenin and vinculin at BCJs in a discontinuous pattern (Figures 4C-D). Staining of tricellulin-RFP with this mechanosensor also showed strong BiFC signals at TCJs, implicating the interaction of α-catenin and vinculin not only at the BCJs but also at the TCJs (Figures 4E).

**Figure 4.**
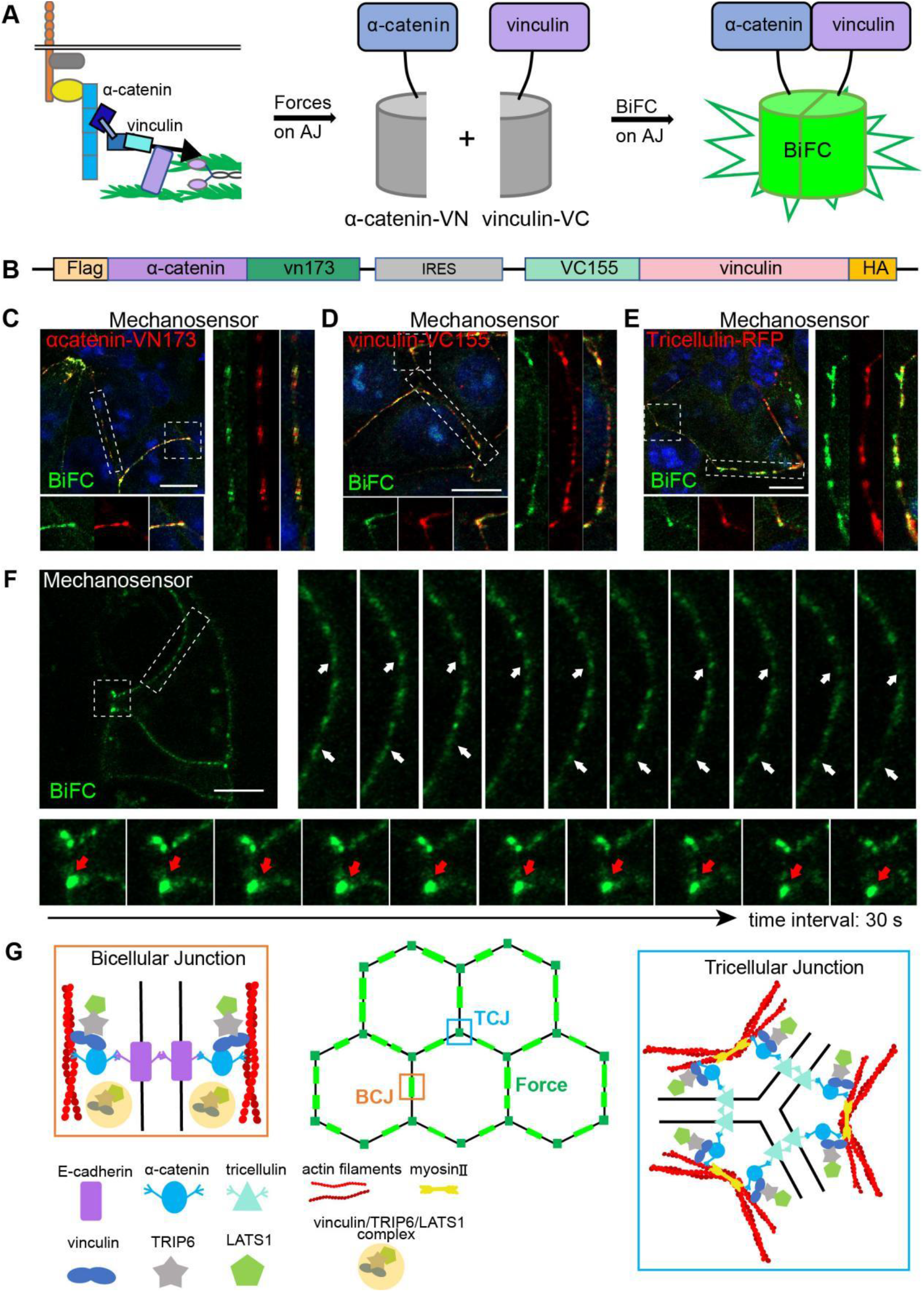
The mechanosensor based on the α-catenin/vinculin complex displays discontinuous and motile signals at BCJs, while stronger signals at TCJs. **A**. Diagram of the mechanosensor design based on the tension dependent interaction between α-catenin and vinculin. **B**. Design of the all-in-one plasmid of the α-catenin and vinculin-involved mechanosensor. Flag-α-catenin-VN173 and VC155-inculin-HA was linked by an IRES sequence. **C-D**. The BiFC signal (green) in the mechanosensor-transfected DLD1 cells, and Flag-α-catenin-VN173 (C) or VC155-inculin-HA (D) were shown in red. **E**. The BiFC signal (green) in the mechanosensor and tricellulin-RFP co-transfected DLD1 cells. **F**. Time-lapse series of DLD1 stable cell lines expressing the mechanosensor based on α-catenin and vinculin interaction. White arrowheads indicate hotspots at BCJs, while red arrowheads indicate the relatively stable BiFC signals at TCJs. Scale bar: 10 μm. **G**. Schematic diagram of the α-catenin/vinculin/TRIP6/LATS1 cassette at the bicellular junctions (BCJs) and tricellular junctions (TCJs). The bright green lines on the hexagon edges show the discontinuous button-like distribution of the tension sensor at BCJs, while dark green dots at the hexagon vertex indicate the stronger localization of the tension sensors at TCJs. All dashed box regions were enlarged on the right. Scale bars are all 10 μm.

We further applied the mechanosensor for live imaging to visualize the forces at AJs. Similar to the staining result, the BiFC signal of the mechanosensor distributed discontinuously at BCJs, and high intensity at TCJs in the live DLD1 cells. The BiFC signals at BCJs appear more dynamic and fluctuating than the signals at TCJs, meaning the forces at BCJs mediated by α-catenin and vinculin are more motile than that of TCJs (Figure 4F, Movies S3 and S4). The time-lapse of BiFC mechanosensor also showed occasionally a BiFC spot at BCJs dynamically moved to and away from a neighboring tension site (Movie S3).

In conclusion, our work demonstrated that the α-catenin/vinculin/TRIP6/LATS1 cassette is a conserved tension sensing complex at both BCJs and TCJs, and uncovered the temporal and spatial characteristics of the junctional tension sensor at BCJs and TCJs. The TRIP6/LATS1 complex forms as puncta in the cytosol together with vinculin but not α-catenin, and were recruited not only to BCJs but also TCJs where α-catenin/vinculin interacts. The tension sensors at BCJs are discontinuously distributed and more motile than TCJs, while are strongly localized at TCJs (Figure 4G).

The α-catenin responds to tension at AJs by changing from a closed self-inhibited configuration to an opened one, thus exposing the vinculin binding site and recruiting vinculin to the sites of tension (Drees et al., 2005; Hirano et al., 2018). The active signals at cell junctions detected by the mechano-split-TurboID, Δmod-Flag as well as the BiFC mechanosensor indicate these are sites where α-catenin interacts with vinculin. α-catenin conformation is found to be kept in an intermediate state under tension (Hoffmann and Dougan, 2012; Maki et al., 2016), Thus, it is possible α-catenin might keep in an unfolded intermediate state in BCJs and TCJs that are under tension. In this study, we did not detect a robust colocalization of YAP and 14-3-3 with α-catenin and vinculin either at BCJs or TCJs, implying that YAP and 14-3-3 are not conservative components of the tension sensor of α-catenin/vinculin complex. TRIP6 and LATS1 were detected both at BCJs and TCJs where α-catenin and vinculin interacts, and appeared as puncta in the cytoplasm with vinculin but not with α-catenin. We speculate that similar to the BCJs, TRIP6 is recruited to TCJs by interacting with vinculin and inhibits LATS1 activity to activate YAP (Dutta et al., 2018).

We also observed α-catenin /vinculin/TRIP6/LATS1 cassette, Δmod-Flag, as well as the mechano-TurboID and BiFC mechanosensor all displayed a discontinuous distribution at BCJs and higher intensity at TCJs, implying that tensions are not evenly exerted on the BCJs. The discontinuous distribution of the tension transducer is more button-like but not zipper-like spots. This might be a way for the cells to protect themselves from being overstretched by the forces at the cell junctions. Whether changes of AJs from a button to a zipper-like pattern, for example in the lymphatics, are accompanied by the discontinuous tension formation still needs to be explored (Baluk and McDonald, 2022). Live imaging further showed the tension sensor at BCJs were more motile compared with TCJs, and dynamically moving to and away from neighbor tension sites. This might be a way to efficiently form a tension transducer complex from an already existing neighbor sites. Application of the BiFC-based mechanosensor in our study to cells and mouse models could be a powerful tool to visualize cell-cell mechanotransduction during tissue development and diseases.

## Methods

### Cell culture and transfection

All cells were cultured with a DMEM medium (Thermo Fisher, US) containing 10% fetal bovine serum (Vivacell, China) and 1% penicillin-streptomycin (Beyotime, China) at 37 °C with 5% CO2. For immunofluorescence staining, DNA transfection of DLD1 cells was performed using the Lipo3000 transfection reagent (Thermo Fisher, US). DNA transfection of HEK293T cells for immunofluorescence staining and immunoprecipitation was performed by the HighGene plus transfection reagent (ABclonal, China). To establish stable α-catenin-VN173-IRES-VC155-vinculin expressing DLD1 cell lines, we infected DLD1 cells with the lentivirus generated from HEK293T cells overnight, followed by selection with 7.5 µg/mL puromycin. HEK293T cells for virus generation were transfected with PEI reagent (Polysciences, US). HEK293T and DLD1 cells were kindly provided by Prof. Jianwei Sun of Yunnan University.

### Immunofluorescence

Cell slides were treated with poly-L-lysine overnight. Slides were then washed with PBS and seeded with cells. Cells were fixed with methanol for 3-5 min at -20°C temperature, then was permeabilized using 0.1% Triton X-100 in PBS for 3 times. Cells were incubated in 1% BSA for 30 min at room temperature, then incubated for 30 min in primary antibody in 1% BSA. After that, cells were washed with 0.1% Triton X-100 in PBS for 3 times, and incubated away from light for 30min in secondary antibody in 1% BSA. Cells were again washed with 0.1% Triton X-100 in PBS for 3 times, and finally were mounted in anti-fluorescence quenching reagents. Antibodies used in the immunofluorescence staining are listed as follows: Monoclonal ANTI-FLAG (Sigma-Aldrich, F1804, 1:1000), HA Tag Monoclonal Antibody (ThermoFisher, 26183, 1:500), Alpha E-Catenin Polyclonal antibody (Proteintech, 12831-1-AP, 1:100), LATS1 Polyclonal antibody (Proteintech, 17049-1-AP, 1:500), TRIP6 Polyclonal antibody (Proteintech, 21163-1-AP, 1:100), DAPI (meilunbio, MA0127, 1:500), CoraLite 488-conjugated Goat AntiMouse IgG (H+L) (Proteintech, SA00013-1, 1:500), CoraLite 488-conjugated Goat AntiRabbit IgG (H+L) (Proteintech, SA00013-2, 1:500), CoraLite 594-conjugated Goat AntiMouse IgG(H+L) (Proteintech, SA00013-3, 1:500), CoraLite 594-conjugated Goat AntiRabbit IgG(H+L) (Proteintech, SA00013-4, 1:500), Alexa Fluor 594 Streptavidin Alexa Fluor (yeasen, 35107ES60, 1:500).

### Western blotting

Cells were lysed using RIPA buffer supplemented with protease inhibitors. Samples collected were separated by SDS-PAGE and transferred onto PVDF membranes. After blocking with 5% nonfat milk at room temperature for 1 hour, the membranes were incubated with primary antibody diluted in 2% nonfat milk for 2.5 hours at RT, followed by washing with 0.1% Tween-20 in PBS for three times. Subsequently, the membranes were incubated with HRP-conjugated secondary antibody diluted with 2% nonfat milk for 1 hour at RT. Finally, the samples were washed again with 0.1% Tween-20 in PBS and imaged using chemiluminescence. Antibodies used in the western blotting are listed as follows: Monoclonal ANTI-FLAG (Sigma-Aldrich, F1804, 1:5000 dilution), HA Tag Monoclonal Antibody (ThermoFisher, 26183, 1:5000), HRP-labeled Streptavidin (Beyotime, A0303, 1:4000), Anti-rabbit IgG, HRP-linked Antibody (Cell Signaling technology, 7074S, 1:4000), Anti-mouse IgG, HRP-linked Antibody (Cell Signaling technology, 7076S, 1:4000).

### Mass Spectrometry Sample Preparation

HEK293T cells were seeded in 10 cm plates. After overnight incubation, cells were transfected with the mechanoTurboID and control plasmids using HighGene reagents, and then treated with biotin (50uM) the next day for 18 hrs. Cells were lysed using RIPA buffer (without SDS and Triton) supplemented with protease inhibitors, and ultrafiltrated using ultrafiltration tube (3 KD, PALL) and washed three times using PBS to exclude free biotin possible detergent in the cell lysates. Then the immunoprecipitation was performed with Streptavidin magnetic beads (Beyotime, China) following the instruction. The purified biotinylated proteins in the beads were sent for mass spectrometry analysis by the Mass Spectrometry Facility of Yunnan University, China following the protocol as described (Geng et al., 2022).

### Plasmids

α-catenin-Flag and α-catenin-mutants-Flag (NM_001323982.2) were constructed by amplification from HaCaT cDNA and cloned into pCMVtag2B using enzyme restriction sites (BamHI or HindIII) by ClonExpress II One-Step Cloning Kit (Vazyme, China). Vinculin (NM_014000.3) was amplified from HaCaT cDNA and cloned into pEGFP-N1 using enzyme restriction sites (HindIII & NotI). LATS1 (NM_004690.4) / YAP (NM_ 001130145.3) / 14-3-3 (NM_006826.4) TRIP6 (NM_003302.3) / Tricellulin (NM_001038603.3) were amplified from HaCaT cDNA respectively and cloned into pEGFP-N1 or pERFP-N1.

### Imaging

For live imaging, glass-bottom dishes were coated with poly-L-lysine for 2 hrs at 37°C, then were washed with PBS and seeded with cells. After overnight incubation, cells were transfected with relative plasmid. After 24h incubation, live cell imaging was performed using a Zeiss LSM800 confocal microscope with Airyscan. Fixed slides were imaged using an inverted Zeiss LSM800 confocal microscope with Airyscan using 63x oil-immersion objective lens and processed using Fiji imageJ.

### Statistics

For the quantification of the colocalization ratio between α-catenin-Flag, α-cateninΔmod-Flag, vinculin-HA and LATS1-RFP with TRIP6-GFP, the number of puncta in the cytoplasm that have colocalization with TRIP6-GFP was numbered and divided by the total number of TRIP6-GFP puncta in the cytoplasm. Data were judged to be statistically significant when p value < 0.05 by one-way ANOVA by Tukey’s multiple comparisons test (ns = not significant, *, p < 0.05; **, p < 0.01, ***, p < 0.001, ****, p < 0.0001). All images were analyzed using ImageJ. Statistical analysis was performed using GraphPad Prism 5 software and Microsoft Excel.

## Acknowledgements

We are grateful to Professor Jianwei Sun from Yunnan Univeristy for the gift of HEK293T cells, Professor Xuna Wu for mass spectrometry analysis, the core facilities of Yunnan University for imaging.

## Fundings

This study was supported by the National Natural Science Foundation of China No. 32270846, Scientific Research Foundation of Education Department of Yunnan Province No. 2023J0003, and Yunnan Fundamental Research Projects No. 202301AU070179.

## Notes

### Competing Interest Statement

The authors have declared no competing interest.

